# Ape cultures do not require behaviour copying

**DOI:** 10.1101/2020.03.25.008177

**Authors:** Alberto Acerbi, William Daniel Snyder, Claudio Tennie

## Abstract

While culture is widespread in the animal kingdom, human culture has been claimed to be special due to being cumulative. It is currently debated which cognitive abilities support cumulative culture, but behavioural form copying is one of the main abilities proposed. One important source of contention is the presence or absence of behaviour copying in our closest living relatives, non-human great apes (apes) – especially given that their behaviour does not show clear signs of cumulation. Those who claim that apes copy behaviour often base this claim on the existence of stable ape cultures in the wild. We developed an individual-based model to test whether ape cultural patterns can emerge in absence of any behaviour copying, when only allowing for a well-supported alternative social learning mechanism, socially mediated reinnovation, where only the frequency of reinnovation is under social influence, but the form of the behaviour is not. Our model reflects wild ape life conditions, including physiological and behavioural needs, demographic and spatial features, and possible genetic and ecological variation between populations. Our results show that, under a wide range of values of parameters, we can reproduce the defining features of wild ape cultural patterns. Overall, our results show that ape cultures can emerge and stabilise without behaviour copying. Ape cultures do not show the signatures of behaviour copying abilities, lending support to the notion that behaviour copying is, among apes, unique in the human lineage. It therefore remains an open question when and why behaviour copying evolved in hominins.

## Introduction

Cumulative culture, the transmission and improvement of knowledge, technologies, and beliefs from individual to individual, and from generation to generation, is key to explain the extraordinary ecological success of our species (1,2). Which cognitive abilities underpin humans’ cumulative cultural capacities, and how these abilities affect the evolution of culture itself are among the most pressing questions of evolutionary human science.

Many species are able to at least use social cues to adjust their behaviour. Various ape species have been shown to posses traditions that are socially influenced in this way (3–7). Humans, by contrast, have cumulative culture. While there are various definitions of cumulative culture (8), some of its characteristics are broadly accepted. Cumulative culture requires the accumulation of cultural traits (more cultural traits are present at generation *g* than at time *g-1*), their improvement (cultural traits at generation *g* are more effective than at generation *g-1*), and ratcheting (the innovation of cultural traits at generation *g* depends on the presence of other traits at generation *g-1*) (9).

While not all human culture needs to be supported by faithful copying (10), cumulative culture depends on an ability to accurately transmit and preserve new behaviours. Experiments have indeed shown that humans are capable of copying behaviours, and that they routinely do so cross-culturally (11,12). More controversial is the claim that other primate species copy behaviours. Arguments regarding the existence of non-human great ape cultures based on behaviour copying raise a puzzling question: if other ape species can and do copy behaviours, why do they not develop cumulative cultures? There are only two possible answers to this question: either apes do not copy behaviour, or copying behaviour does not automatically lead to cumulative culture.

Primatologists have claimed the existence of ape cultures based on the ability of behavioural forms copying, drawing on observations conducted on wild ape populations. For example, researchers examined the population-level distribution of behaviours in populations of chimpanzees across seven sites, and argued that the inter-site differences in the frequency of behaviours proved the existence of behaviour copying-based cultures in these populations (3). We developed an individual-based model to assess whether these patterns, and similar patterns found in orangutans (5), spider monkeys (13), gorillas (6), and bonobos (14), actually justify the conclusion that behaviour copying is the underlying learning mechanism. We focus, as illustration, on the original study (3), but the model is aimed to illustrate, more generally, how non-copying mechanisms can generate population-level distributions of behavioural traits that have been interpreted as proof of individual-level copying abilities.

While our hypothetical ape species, “oranzees”, can be influenced by social cues (widespread in the animal kingdom, and certainly present in all apes), we did not implement behaviour copying. More specifically, oranzees acquire behaviours through socially mediated reinnovation (15): all behavioural forms are already latent in the behavioural repertoire of oranzees, meaning that these form can be reconstructed (“innovated”) by individual learning. Initial innovation, by acting as cues, can trigger reinnovations in others: the behavioural forms are not under social influence, but their freuqency is. Our results show that, under realistic values of the main parameters, this mechanism can reproduce the distribution of behavioural traits found in (3). In other words, as oranzees can and do show cultural patterns resembling wild ape patterns, this shows that such patterns do not allow to conclude that behaviour copying must have taken place.

## Materials and methods

We build an individual-based model that reproduces a world inhabited by six populations of “oranzees”, a hypothetical ape species. The model is spatially explicit: the oranzees populations are located at relative positions analogous to the six chimpanzees sites in (3). This is important to determine the potential genetic predispositions and ecological availabilities associated with their possible behaviours (see below). Population sizes are also taken from the sites in (3). Following (16), we use data from (17), and we define population sizes as *N* = {20; 42; 49; 76; 50; 95}.

Oranzees are subject to an age-dependent birth/death process, broadly inspired by descriptions in (18). A time step *t* of the simulation represents a month in oranzees’ life. From when they are 25 years old (*t* = 300), there is a 1% probability an oranzee will die each month (maximum lifetime is capped at 60 years, i.e. *t* = 720). The number of individuals in the population is fixed, so each time an oranzee dies it is replaced by a newborn.

A newborn oranzee does not yet show any behaviour, but is individually capable of developing them. Behaviours can be developed at each time step, among the 64 possible behaviours. The process of development of behaviours is influenced by: (i) the oranzees ‘state’, which depends on the behaviours an individual already expresses, (ii) the frequency of the behaviours already expressed in the population (“socially mediated reinnovation”), and (iii) the genetic propensity and ecological availability locally associated with the behaviour. At the beginning of the simulations, the populations are randomly initialized with individuals between 0 and 25 years old.

### Oranzees’ behaviours and state

In the oranzees’ world, 64 behavioural form are latently possible (loosely modelled on the 65 behaviours coded in (3), but making it an even number from modelling convenience). Behaviours are divided into two categories: 32 social and 32 food-related behaviours. These figures where chosen to resemble the behavioural categories considered in (3). Behaviours serve oranzees to fulfill various goals. Oranzees have a ‘state’ that is based on how many goals are fulfilled in the two main categories of social and food-related behaviours.

In the case of social behaviours, we further assume four sub-categories (‘play’, ‘display’, ‘groom’, ‘courtship’; note tht the names are only evocative), each with eight possible different behaviours that serve the same goal. A goal is considered fulfilled if an oranzee shows at least one behaviour out of the eight in the sub-category. Oranzees have a ‘state’ that is based on how many of the four goals are fulfilled. An oranzee has a state value of 0.25 if, for example, it shows at least one behaviour in the category ‘play’, and none of the others, and a state value of 1 if it shows at least one behaviour in each sub-category. *p*_social_, the probability to innovate a social behaviour, is drawn from a normal distribution with mean equal to 1 – *state*_social_.

Food-related behaviours are analogously divided into sub-categories. Differently from social behaviours, there is a variable number of behaviours in each sub-category. In addition, sub-categories are associated to two different ‘nutrients’, *Y* and *Z*. Here individuals need to balance their nutritional intake, so that their optimal diet consist in a roughly equal number of food for one and the other nutrient. The state, for food-related behaviours, depends on the total amount of food ingested *and* on the balance between nutrients. The state is calculated as the sum of each sub-category fulfilled (as above, for this to happen there needs to be at least one behaviour present) minus the difference between the number of sub-categories providing nutrient *Y* and the number of sub-categories providing nutrient *Z*. We normalize the state between 0 and 1, and, as above, *p*_food_ is then calculated as 1 – *state*_food_.

### Socially mediated reinnovation

At each time step, all oranzees have a probability of individual innovation for social and food-related behavioural forms calculated as described above. The specific behavioural form an oranzee will acquire depends both on the frequency of the behaviours that are already present in the population (see below), and on the ecological availability and genetic propensity associated to the behavioural form. A further parameter of the model, *S*, controls the probability that each reinnovation is socially mediated (15). When a reinnovation is socially mediated, the probability of innovating each behaviour *B_i_* is weighted by its proportional instances in the population among the behaviours of the same category (social or food-related). That is, the frequency of behavioural forms can catalyse more individual innovations of the same behaviour: common behaviours are more likely to be individually reinnovated.

When the innovation is not socially mediated, the probability of innovating each behaviour is random. Only one behaviour per category can be innovated at each time step.

### Genetic propensity and ecological availability

The behaviour selected in the previous step is then innovated or not according to its genetic propensity and, in case of food-related behaviours, ecological availability.

Genetic propensity is a probability *p_g_*(0, 1), assigned independently for each of the 64 behaviours. A parameter of the model, *α_g_*, determines the probability that the genetic propensity of each behaviour is equal for all the six populations or whether is different. If the probability is equal, *p_g_* is randomly drawn. If it is different, we assign the propensity using a geographical gradient. We choose a random point and calculate its distance to each population. Distances are then transformed to *p_g_* by rescaling them between 0 and 1, so that for the farthest site where *p_g_* = 0, the associated behaviour cannot possibly be expressed (see SI). Notice that *α_g_* = 0 does not mean that there are no genetic influences on the behaviour, but that there are no *differences* between the populations with regard to this aspect.

Ecological availability is a probability *p_e_*(0, 1) that represents the likelihood of finding a resource, or its nutritional value, in each site. Ecological availability is assigned only to food-related behaviours, and it is calculated in the same way of *p_g_*, using the parameter *α_e_* to determine the probability of ecological availability being different in the six populations.

### Model’s output

We run simulations for *t*_max_ = 6000 (corresponding to 500 years of oranzee-time). For each simulation, following (3), we classify each behaviour, in each population, as:

- *customary*: a behaviour observed in over 50% of individuals in at least one age class (see SI for how age classes are defined in our model).
- *habitual*: a behaviour observed in at least two individuals across the population.
- *present*: a behaviour observed in at least one individual across the population.
- *absent*: a behaviour not observed even once in the population.
- *ecological explanations:* a behaviour that is absent due to a complete lack of local ecological availability (i.e., in our model, associated to *p_e_* = 0).

Notice that one category in (3) (*unknown*, i.e. “the behaviour has not been recorded, but this may be due to inadequacy of relevant observational opportunities”) does not apply in our case, because we have complete knowledge of the output of the simulations.

Finally, to test how well our model compares to wild apes, we calculate the same “patterns” described in (3):

- *A*: behaviour absent at no site.
- *B*: behaviour not achieving habitual frequencies at any site.
- *C*: behaviour for which any absence can be explained by local ecological factors.
- *D*: behaviour customary or habitual at some sites yet absent at others, with no ecological explanation, i.e. behaviours defined as “cultural”.

Further details of the model implementation and of how outputs are processed are available in SI. The full code of the model allowing to reproduce all our results, plus a detailed description of the model development is available in a dedicated GitHub repository, at https://github.com/albertoacerbi/oranzees.

## Results

We are particularly interested in the realistic parameter conditions of moderate to high environmental variability (i.e. *α_e_* from 0.5 to 1) and zero to moderate genetic differences (i.e. *α_g_* from 0 to 0.5). We ran 20 simulations for each combination (for a total of 600 runs). For all, reinnovation is socially mediated (*S* = 1). The results show that various combinations of parameters produce a number of cultural behaviours (pattern *D*) consistent with the general pattern described in (3), in absence of any explicit copying mechanism being implemented (see Figure 1). In Figure 2, we reproduce the output of a run where 38 cultural behaviours were found, and how they were classified in each of the six simulated populations, using a visualization inspired by (3).

**Figure 1:**
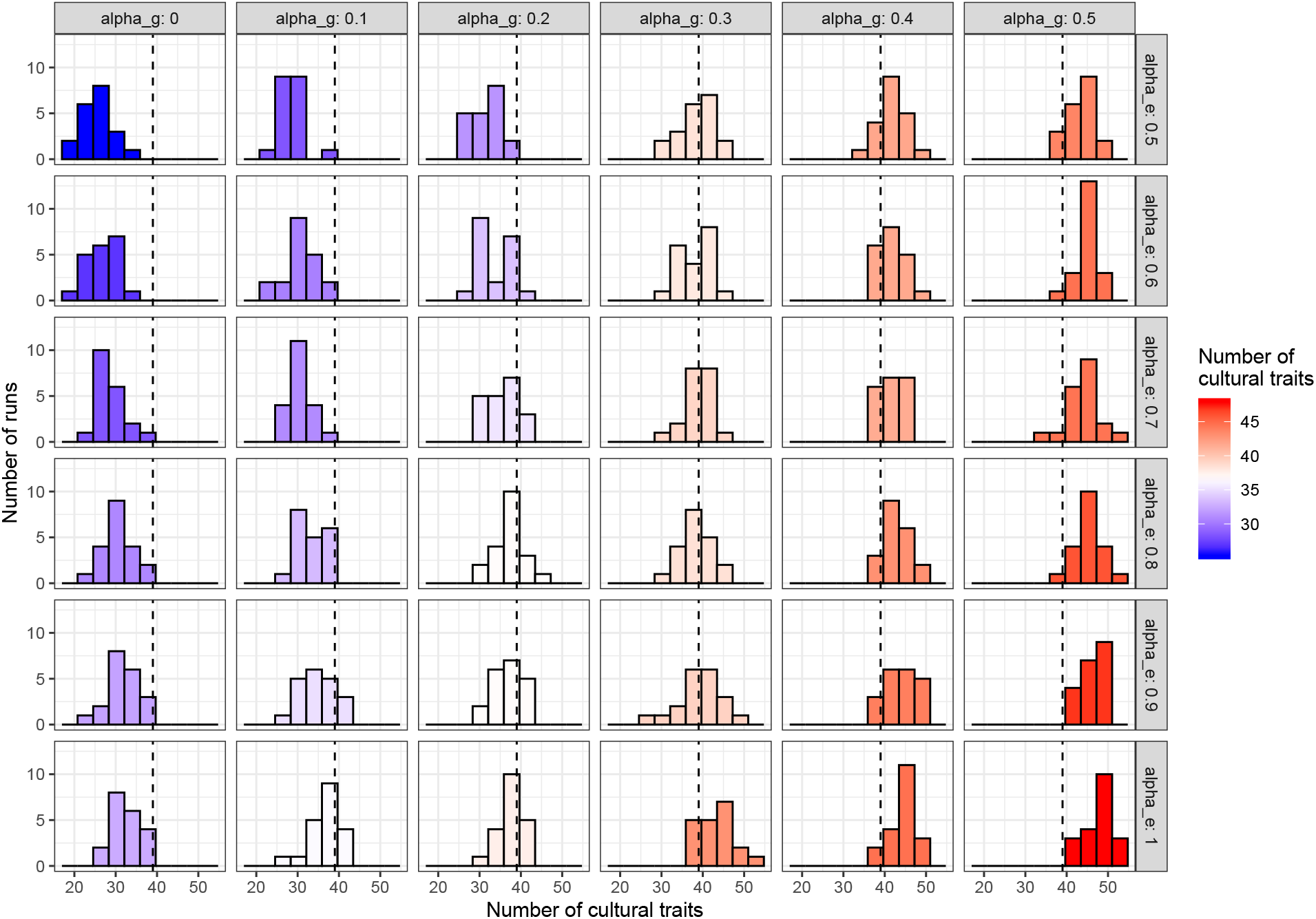
Number of cultural traits in oranzees, when varying ecological and genetic diversity. Red color indicates simulation runs that produced more than 38 cultural traits (the number of cultural traits identified in 1); blue color indicates simulation runs that produced less than 38 cultural traits. For all simulations, *S* = 1, *α_e_* and *α_g_* as indicated in the plot. *N* = 20 runs for each parameters combination. (See SI for other values of *S*, *α_e_*, and *α_g_*, including all equal to zero.)

**Figure 2:**
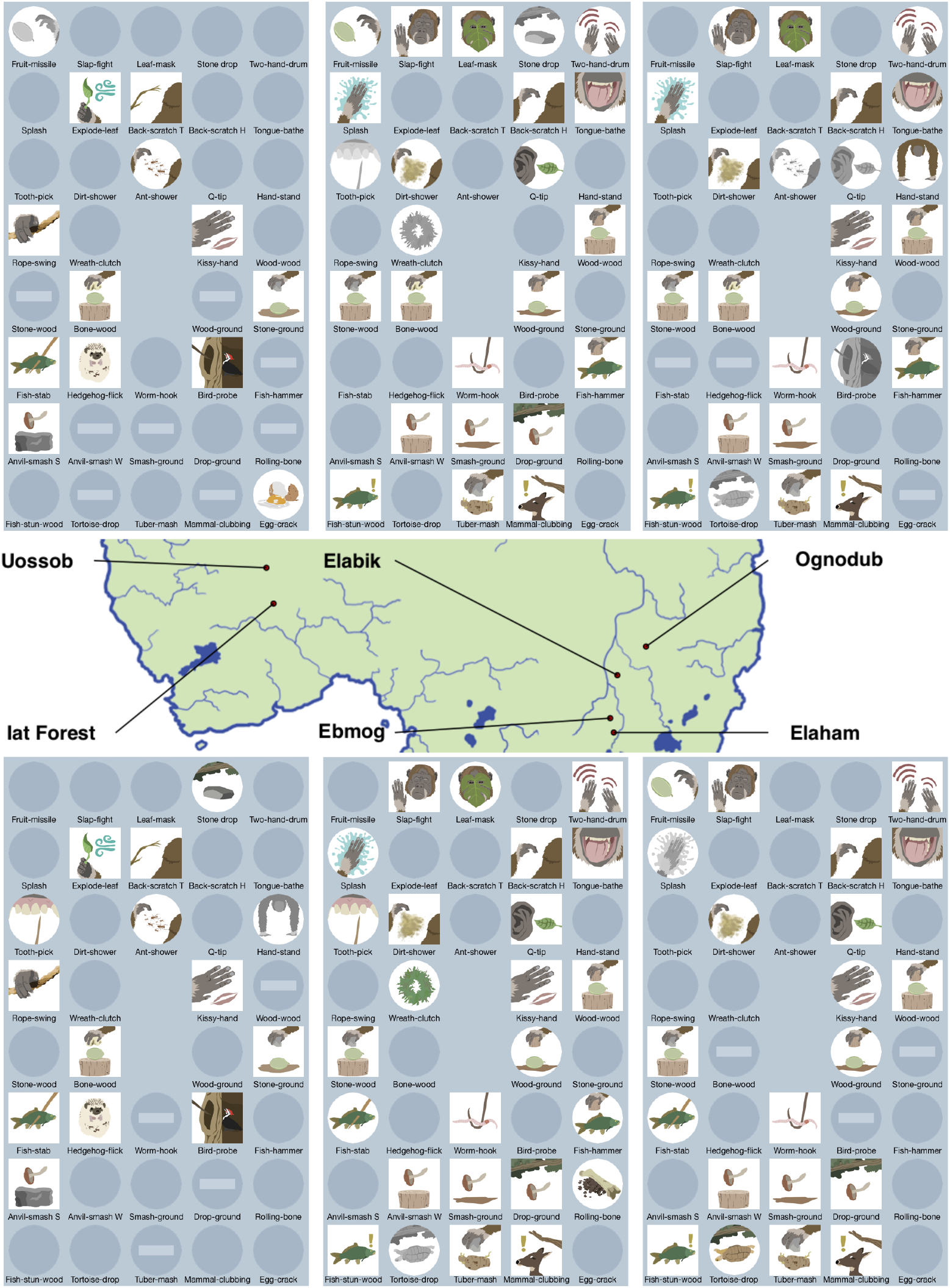
Example of a simulation run that produces 38 cultural traits (*S* =1, *α_e_* = 0.8, and *α_g_* = 0.2). Color icons indicate customary behaviours; circular icons, habitual; monochrome icons, present; clear, absent; horizontal bar, absent with ecological explanation. The names of the behaviours are only evocative, see SI for a complete list.

We also analysed the effect of the parameter *S* (proportion of socially mediated reinnovations), in three conditions (see Figure S4): (a) no genetic differences and intermediate ecological differences (compare to the high-left corner of Figure 1, where with *S* = 1 simulations produce less than 38 cultural behaviours), (b) one of the conditions that produce good match with (3), namely *α_e_* = 0.8 and *α_g_* = 0.2, and (c) intermediate genetic differences and high ecological differences (compare to the low-right corner of Figure 1, where with *S* = 1 simulations produce more than 38 cultural behaviours). As expected, decreasing *S* decreases the number of cultural behaviours. Conditions where, with *S* = 1, there were more than 38 cultural behaviours could still produce results analogous to (3), given that not all reinnovations are socially mediated.

As a further proof of our model’s fit with empirical data, our outputs not only accurately reproduce the number of cultural behaviours (pattern *D*), but also the number of behaviours classified in the other three patterns (*A*, *B*, *C*, see above) in (3) (see Figure S5).

Finally, we ran 100 simulations for one of the conditions where we have a good match for the number of cultural behaviours in (3) (*α_e_* = 0.8; *α_g_* = 0.2, *S* = 1). In each simulation, we recorded, for each population, the number of behaviours (habitual + customary + present) that are also classified as cultural (see Figure S6). We find a small, but significant, correlation between population size and number of cultural traits (*p* < 0.00001, *ρ* = 0.2, *N* = 600). In other words, our model reproduces an effect of cultural accumulation (i.e. increased number of expressed behaviours) relative to population size possibly found in real populations - see (16,19,20) - again, in the absence of behaviour copying.

## Discussion

We developed an individual-based model to examine under which conditions a distribution of cultural traits analogous to the distribution reported in (3), a representative study of primate culture, could emerge, crucially, without implementing any behaviour copying mechanism. We modelled various details of the original wild ape study, including demographic and spatial features, as well as effects of genetic propensity and ecological availability on the behaviours. Given the widespread availability of non-copying variants of social learning across the animal kingdom, we also included socially mediated reinnovation, where social learning merely catalyses individual reinnovation, without any behavioural form copying (15).

It is important to notice that our model do not, and cannot, *exclude* that the behavioural distributions observed in wild apes are produced by copying mechanisms at the individual level. What our model does, however, is showing that behaviour copying is not necessary and other mechanisms are sufficient to generate analogous distributions.

Socially mediated reinnovation is implemented in the model as a sampling biased by the frequency of the observed behavioural forms. The fact that this may be interpreted as equivalent to copying makes exactly our point.

The model does not allow for the specifics of behavioural forms to be transmitted (in contrast to, e.g. (21)) Instead, the presence of behavioural form merely acts as a trigger, creating an illusion of behaviour copying (in humans, this happens for example during contagious yawning - where yawning likewise acts as a trigger only, and the form of yawns themselves are not copied). Given the empirical support for the existence of socially mediated reinnovation in apes (22,23) and the absence of spontaneous copying of behavioural forms in apes (24,25), our model’s most parsimonious interpretation has to be that wild ape cultural patterns do not allow to conclude for the presence of behaviour copying.

Our main result is that we can reproduce the general pattern observed in populations of wild apes under realistic values of the parameters of genetic propensity and ecological availability, namely zero to medium importance of genetic variation, and medium to high importance of ecological variation. (Notice however that even in the entire absence of any ecological and genetic variation, i.e. with *α_e_* = 0 and *α_g_* = 0, some cultural traits occur, see Figure S7). More generally, the model shows that behaviour copying is not logically required for patterns interpreted as cultural in primates studies (3,5–7,13,14) to emerge. In addition, and as further support for our results, our model not only reproduces the ape cultural behavioural patterns, but also the proportions among the other patterns observed in wild apes, i.e. absent behaviours, behaviours not achieving habitual frequencies at any site, and behaviours absent because of ecological factors. The exact number of behaviours for each patterns depend on model parameters, including the choice of considering a total set of 64 possible latent behaviours that can vary between populations. The general conclusion that cultural patterns can be generated hold regardless the exact number of behaviour considered and, given that (3) selected this number based on relatively informal criteria, we used a similar strategy here.

In our model, we focused on the mechanism of socially mediated reinnovation, that is, we assumed that members of our hypothetical species, oranzees, had a probability to individually reinnovate the details of a specific behavioural form stochastically linked to how many other oranzees in the population were already showing this behaviour. Mere aspects of these behaviours (e.g. presence of sticks near prey) act as cues that trigger individual renovation of behavioural forms in others. While this is a realistic assumption (23) and while it reproduces in our model the chimpanzees’ cultural pattern observed in realistic conditions, our results demonstrate that even this is not always necessary. Given certain combinations of parameters, such as higher genetic and ecological diversities, analogous population level patterns can be obtained even when reinnovation is not socially mediated, i.e. when oranzees are not influenced by the behaviours of the other individuals in their populations (compare figure S4). That is, similar patterns can exist when the underpinning individual-level mechanisms are not cultural even in a minimal sense (26). However, socially mediated reinnovation is likely required to explain observed differences in behavioural frequencies between the subset of ape populations that exist in genetic contact and that share similar environments (27).

Finally, our model reproduces a reported correlation between population size and number of cultural traits in the six populations (16,19,20). The magnitude of the effect is small, which is to be expected, given that the presence of this correlation in real populations of (human and non-human) apes is currently debated (28). Notice that this correlation too is brought about without any behaviour copying, so that there is no need to invoke reasons concerning details of cultural transmssion (e.g. (29)) to explain such a pattern.

More generally, the results of our models suggest caution when deriving individual-level mechanisms from population-level patterns (see also (30,31)). Cultural systems, as many others, exhibit equifinality: the same global state can be produced by different local underlying processes. Models and experiments are crucial to test the plausibility of inferences going from global to local properties.

In conclusion, our model strongly suggests that the data available on the behavioural distributions of apes populations cannot demonstrate that ape possess cultures influenced by behaviour copying, let alone *requiring* behaviour copying. This, in turn, may provide an explanation to why ape cultures are not cumulative: if cumulative culture requires at minimum behavioural form copying, we should not expect any species lacking this mechanism to produce and maintain cumulative culture. Given the phylogenetic closeness of apes to the human lineage, our results speak also of the likely absence of behaviour copying of the last common ancestor of apes and humans (32).

## Acknowledgements

This project has received funding from the European Research Council (ERC) under the European Union’s Horizon 2020 research and innovation programme (grant agreement n° 714658; STONECULT project). We would like to than Mima Batalovic for the support provided, and Elisa Bandini, Alex Mielke, Alba Motes Rodrigo, and Jonathan Reeves for comments on earlier versions of the manuscript.

