## Supplementary Information for "Ape cultures do not require behaviour copying"

#### Additional model information

This document provides additional information on the individual-based model described in the paper *Ape cultures do not require behaviour copying*. The full code to run the model and to reproduce the results can be found in https://github.com/albertoacerbi/oranzees, together with a detailed documentation of the model development.

##### The oranzees' world

The oranzees model is an individual-based model, fully written in R, that reproduces a world where six populations of "oranzees" (a hypothetical ape species) live. The model is spatially-explicit: the six populations are located at relative positions analogous to the six populations of chimpanzees in (1) - see Figure 1. For modelling convenience, we put these locations approximately in the center of a 1000 x 1000 squared environment in order to be able to process their relative distances, that we use to calculate genetic propensity and ecological availability of the behaviours (see below).

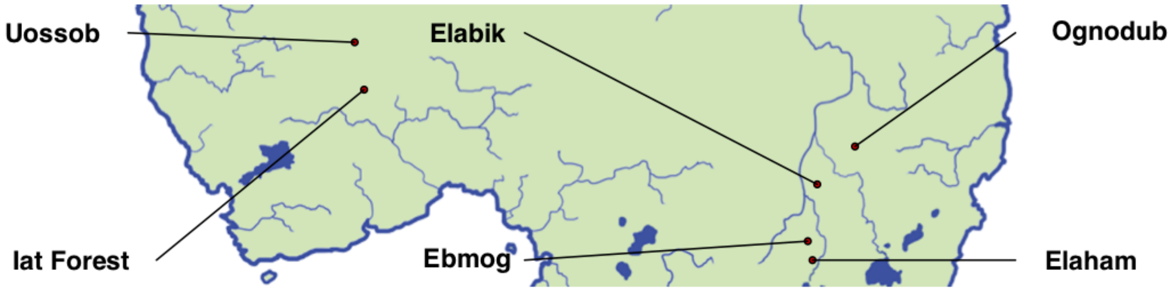


Figure 1: Location of the six populations of oranzees.

The population sizes are also modelled on the real chimpanzees populations considered in the study above. Following (2), we use data from (3):

| Group | Population size |
|---|---|
| Uossob | 20 |
| Elabik | 42 |
| Ognodub | 49 |
| Iat Forest | 76 |
| Ebmog | 50 |
| Elaham | 95 |

#### Geographical gradient for genetic propensity and ecological availability

As described in the main manuscript, two parameters of the models, $\alpha_g$ and $\alpha_e$, determine the probability that the genetic propensity and ecological availability associated to the behaviours are equal for all the six populations, or if they differ among the populations.

Independently for each behaviour, if genetic propensity (or ecological availability) is equal, the probability associated ($p_g$ or $p_e$) is a randomly drawn number between 0 and 1, the same for all six populations. If they are not equal, the values of $p_g$ (or $p_e$) are assigned using a geographical gradient, by choosing a random point in the oranzees' world, and calculating its distance to each population. Distances are then transformed to $p_g$ (or $p_e$) by rescaling them between 0 and 1, so that for the farther population $p_g = 0$ i.e. the associated behaviour will be impossible to express (or $p_e = 0$ i.e. the associated behaviour will be absent with an "ecological explanation").

In the example in Figure 2, a particular behaviour will have $p_g = 1$ (or $p_e = 1$) in the Ognodub site, $p_g = 0$ (or $p_e = 0$) in Iat Forest and Uossob, and intermediate values in the other sites.

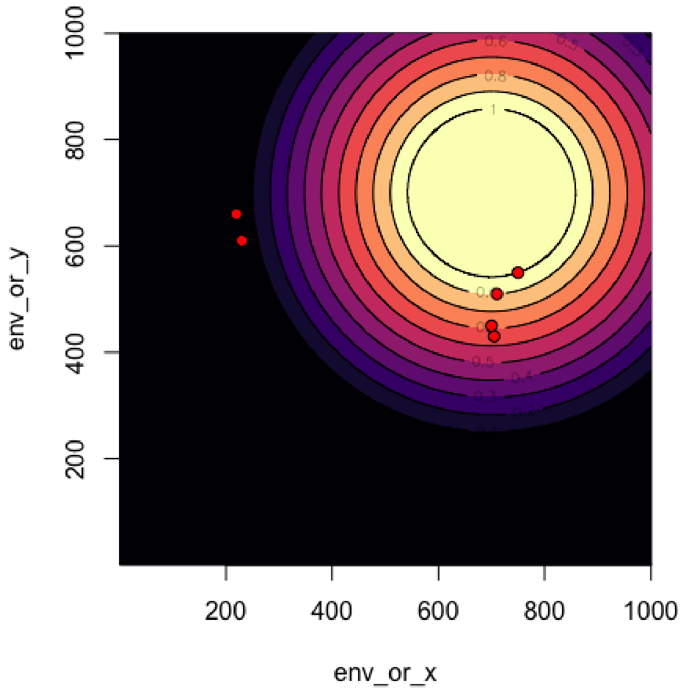


Figure 2: Example of calcualtion of $p_g$ (or $p_e$). The red points represent the oranzee populations. The color gradient represents the value of $p_g$ (or $p_e$).

#### Sub-categories of behaviours

There are 64 behaviours possible in the model (inspired by the 65 coded in (1), but made into an even number for modelling convenience), divided in two main categories: "social" and "food-related". Each category is further subdivided in sub-categories. Sub-categories, for food-related behaviour, are further assigned to specific "nutrients". This information is used to calculate the oranzee's state according to its behaviour (see main manuscript). The names of behaviours and of the sub-categories are only evocative. They are used to illustrate our results in Figure 2 (main manuscript).

##### Social

| Sub-category | behaviour |
|---|---|
| Play | fruit-missile |
| Play | slap-fight |
| Play | air-split |
| Play | leaf-mask |
| Play | whistle |
| Play | pebble-tease |
| Play | tumbling |
| Play | brick-fall |
| Display | stone drop |
| Display | branch pull-release |
| Display | arm-cross |
| Display | two-hand-drum |
| Display | splash |
| Display | arm-swing |
| Display | explode-leaf |
| Display | contorsionist |
| Groom | tool back-scratcher |
| Groom | hand back-scratcher |
| Groom | tongue-bathe |
| Groom | tooth-pick |
| Groom | dirt-shower |
| Groom | ant-shower |
| Groom | q-tip |
| Groom | exfoliate-fruit |
| Courtship | flower-offer |
| Courtship | hand-stand |
| Courtship | rope-swing |
| Courtship | leaf-fan |
| Courtship | wreath-clutch |
| Courtship | ear-pull |
| Courtship | kissy-hand |
| Courtship | hop-dance |

**Food-related**

| Sub-category | behaviour | Nutrient |
|---|---|---|
| Fruit-hammer foraging | wood-wood | Y |
| Fruit-hammer foraging | wood-stone | Y |
| Fruit-hammer foraging | stone-wood | Y |
| Fruit-hammer foraging | stone-stone | Y |
| Fruit-hammer foraging | bone-wood | Y |
| Fruit-hammer foraging | bone-stone | Y |
| Fruit-hammer foraging | wood-ground | Y |
| Fruit-hammer foraging | stone-ground | Y |
| Stick-based foraging | stick-throw V | Z |
| Stick-based foraging | stick-throw A | Z |
| Stick-based foraging | fish-stab | Z |
| Stick-based foraging | hedgehog-flick | Z |
| Stick-based foraging | worm-hook | Z |
| Stick-based foraging | bird-probe | Z |
| Stick-based foraging | fish-hammer | Z |
| Stick-based foraging | spin-seed | Z |
| Anvil smash | anvil-smash S | Y |
| Anvil smash | anvil-smash W | Y |
| Anvil smash | smash-ground | Y |
| Anvil smash | drop-ground | Y |
| Rolling pin techniques | rolling-wood | Z |
| Rolling pin techniques | rolling-stone | Z |
| Rolling pin techniques | rolling-bone | Z |
| Rolling pin techniques | rolling-other | Z |
| Insect swatting | bug-clap | Y |
| Insect swatting | stick-insect | Y |
| Fish stunning | fish-stun-stone | Z |
| Fish stunning | fish-stun-wood | Z |
| Tortoise-flip | tortoise-drop-on-stone | Y |
| Potato-mash | tuber-mash | Z |
| Clubbing | mammal-clubbing | Y |
| Egg cracking | egg-crack | Z |

#### Example of single run

Figure 3 shows an example of the entire history of all behaviours in a single run, for a single population (geographical location and population size are based on "Uossob"), with $\alpha_g = 0.2$, $\alpha_e = 0.8$, and $S = 1$, i.e. one of the combinations of parameters that produces a number of cultural behaviours similar to (1).

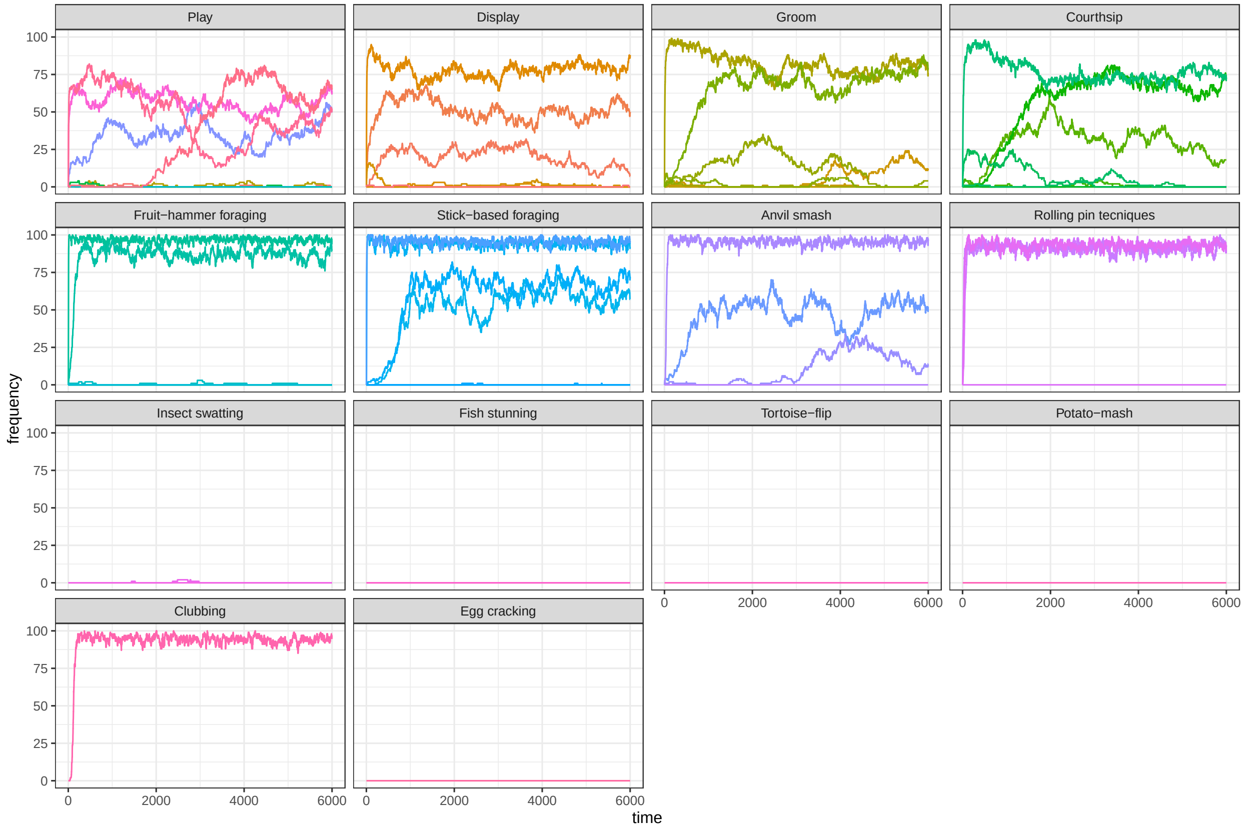


Figure 3: Example of a single run with $\alpha_g = 0.2$, $\alpha_e = 0.8$, and $S = 1$. The plots show the frequencies of the 64 possible behaviours, divided in panels by sub-category.

#### Age classes to calculate customary behaviours

To determine *customary* behaviours, we need to define age classes for individuals (the definition of customary behaviours, from (1) is a behaviour observed in over 50% of individuals in at least one age class). We define three age classes as follows:

- *adults*: individuals that are more than 16 years old.
- *subadults*: individuals between 8 and 16 years old.
- *juveniles*: individuals that are less than 8 years old.

### Supplementary figures

#### Figure S4

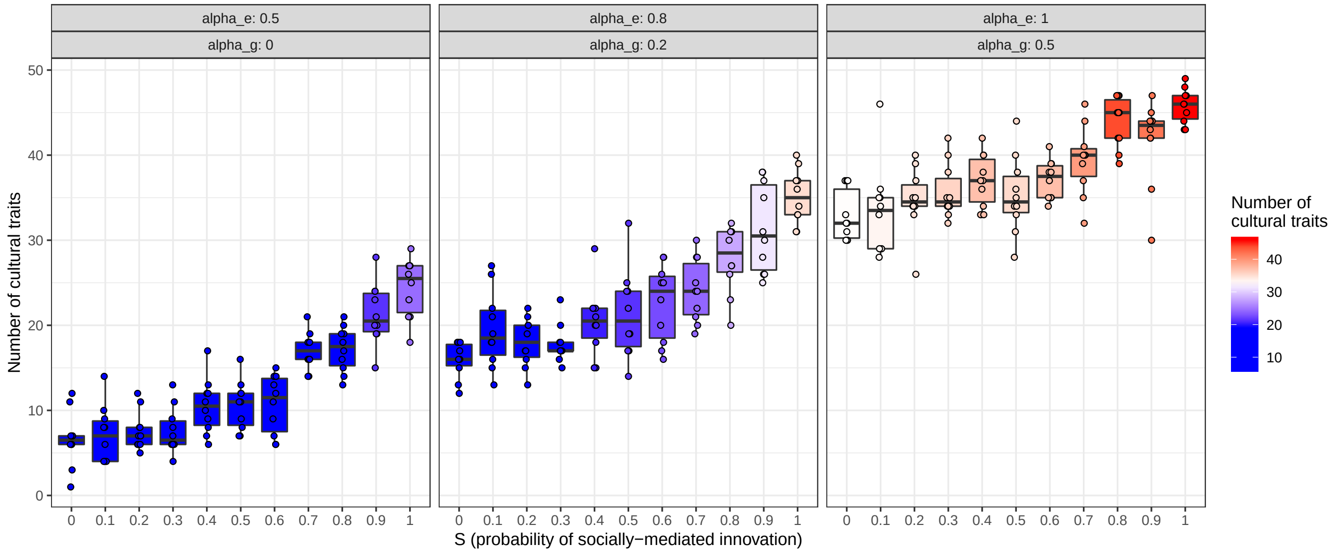


Figure 4: Cultural traits in oranzees, after varying the probability of socially-mediated innovations. Red color indicates simulation runs that produced more than 38 cultural behaviours (the number of cultural traits identified in 2); blue color indicates simulation runs that produces less than 38 cultural behaviours. $S$, $\alpha_e$ and $\alpha_g$ as indicated in the plot. $N = 10$ runs for each combination of parameters.

#### Figure S5

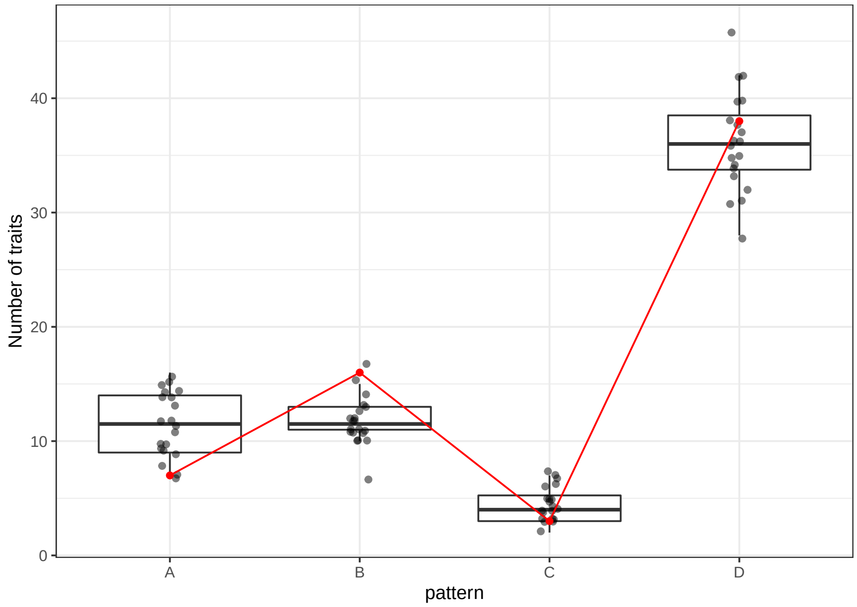


Figure 5: Number of traits for each of the four patterns (*A*, *B*, *C*, *D*, see description in the main manuscript) for the parameters $\alpha_e = 0.8; \alpha_g = 0.2, S = 1$. The red values are the values described for real chimpanzees populations. $N = 20$ runs.

#### Figure S6

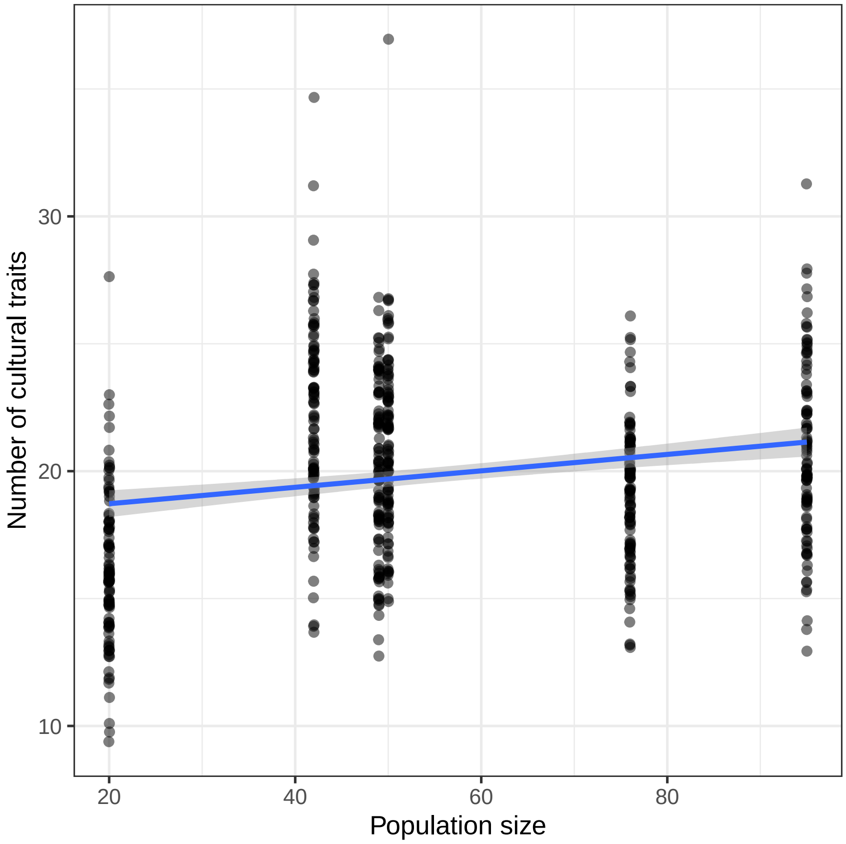


Figure 6: Number of cultural traits for each population for the parameters $\alpha_e = 0.8; \alpha_g = 0.2, S = 1$. The blue line is a linear fit of the data. $N = 100$ runs.

**Figure S7**

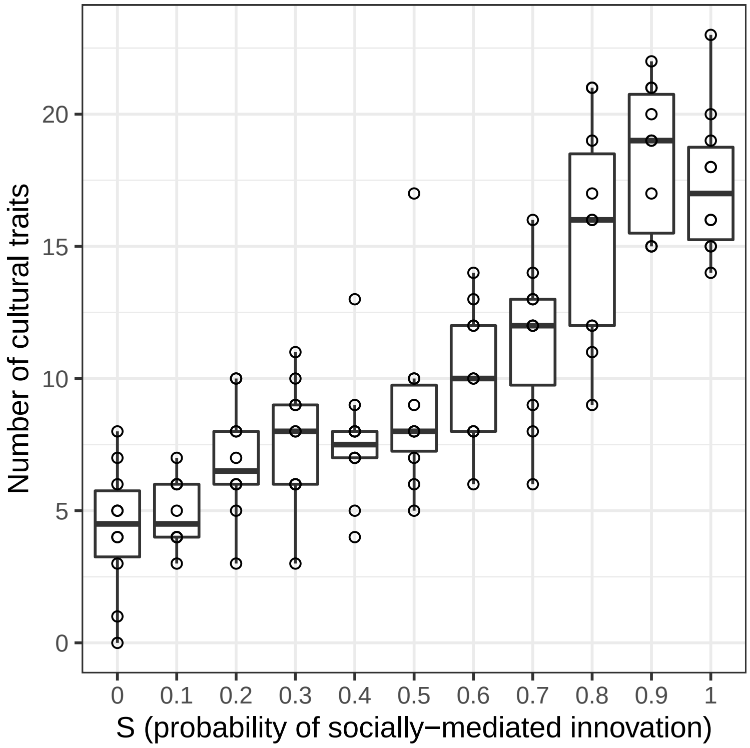


Figure 7: Number of cultural traits in complete absence of ecological and genetic variation ($\alpha_e = 0; \alpha_g = 0$), woth varying probability of socially-mediated innovations ($S$). $N = 10$ runs for each value of $S$. Note that culture-like patterns often remain even in the total absence of genetic differences, environmental differences and social learning (result on far left). To the best of our knowledge this is a result not previously described in the literature - though we suspect these cultures would not be stable over time.
